# Engineering light robustness: Adaptive evolution uncovers new genetic determinants of HL tolerance in *Synechocystis*

**DOI:** 10.1101/2025.07.27.667079

**Authors:** Tao Sun, Kungang Pan, Yaru Xie, Shubin Li, Congzhuang Li, Dailin Liu, Xiaofei Zhu, Weiwen Zhang, Lei Chen

## Abstract

Excess light absorption causes severe photo-oxidative damage and limits photosynthetic efficiency. Enhancing high-light (HL) tolerance and energy utilization in photosynthetic cyanobacteria holds great potential for sustainable biomanufacturing. Here, using *Synechocystis* sp. PCC 6803 (Syn6803) as a model, we generated eight independently evolved strains tolerant to hyper-HL (2000 μmol photons/m^2^/s) through approximately two years of natural adaptive laboratory evolution (ALE). Remarkably, these strains circumvented common evolutionary trade-offs; all demonstrated markedly enhanced quantum yields and photosynthetic performance, with four strains exhibiting a 121.71%-168.36% increase in biomass accumulation under HL. When equipped with the heterologous sucrose transporter CscB, sucrose productivity under HL increased by up to 208.33%. Whole-genome resequencing unveiled a total of 77 mutations across the eight lineages, uncovering a diverse polygenic landscape for HL adaptation. Reverse genetics combined with structural modeling highlighted pivotal roles for regulatory and structural loci, including *slr0758* (KaiC1) and *slr1513* (SbtB), in modulating circadian rhythmicity and inorganic carbon uptake to prevent photo-oxidative overload. Overall, our findings uncover diverse evolutionary routes toward high-light acclimation and provide superior cyanobacterial chassis for photosynthetic biotechnology.

## Introduction

Photosynthesis is the most important chemical reaction on earth, which contributes to the fixation and conversion of hundreds of billion tons of CO_2_ annually^1^. Nevertheless, photodamage and photoinhibition caused by high light (HL) can lead to the over accumulation of reactive oxygen species (ROS) in photosynthetic organisms, disrupting their photosystems and key cellular components like nucleic acids, lipids and proteins thus reducing the global primary productivity potential^2^. Cyanobacteria are the only prokaryotes performing oxygenic photosynthesis, which account for more than 20% of global carbon fixation^3^. Given their relatively faster growth rate, uncomplex genetic makeup and well-established genetic toolkits ^4^, cyanobacteria have been considered an ideal model for studies on photosynthesis. Meanwhile, “photoautotrophic cell factories” based on engineered cyanobacteria have realized the production of dozens of different chemicals and biofuels^5, 6^. Therefore, enhancing the tolerance of cyanobacteria to HL is not only beneficial for creating robust chassis but can also guide the modification of plants, especially since cyanobacteria are thought to be the ancestors of chloroplasts^7^.

Traditionally, besides the accelerated turnover of photodamaged PSII, cyanobacteria have evolved multiple strategies to cope with high light stress, including reducing the harvest of light energy, quenching excess light energy and inducing alternative electron transport pathways^2, 8, 9^. In response to high light stress, cyanobacteria tend to decrease the size of the harvest antenna phycobilisomes (PB) and the amount of chlorophyll on reaction centers to avoid excess light absorption^10^. Additionally, a special nonphotochemical quenching (NPQ) process mediated by the orange carotenoid protein (OCP) in cyanobacteria helps dissipate energy into heat under HL conditions^11, 12^. Furthermore, cyanobacteria can utilize the photorespiratory pathway to consume extra ATP and NADPH by converting 2-phosphoglycolate into CO_2_, NH_4_^+^, and 3-phosphoglycolate^13^. However, these mechanisms are ineffective when exposed to light densities beyond their tolerance, such as 200 μmol photons/m^2^/s for *Synechocystis* sp. PCC 6803 (hereafter Syn6803) and 500 μmol photons/m^2^/s for *Synechococcus elongatus* PCC 7942.^9, 14^ (It is worth noting that the actual light tolerance for cyanobacteria may vary among different studies due to differences in light source, spectrum, CO_2_ concentrations, and cultivation temperature). Therefore, there is still a need for robust cyanobacterial cells that can withstand HL.

Previously, two pioneering studies conducted by Dann et al.^15^ and Yoshikawa et al.^16^ reported the generation of HL-tolerant Syn6803 via chemical mutagen-aided adaptive laboratory evolution (ALE) and short ALE, respectively, leading to the identification of genes including *slr0844, sll1098, hik26*, and *slr1916* responsible for HL tolerance. Here, through approximately two years of natural ALE without chemical mutagens, we generated eight independent Syn6803 lineages capable of robust growth under extreme high light (2000 μmol photons/m^2^/s). Unlike previously reported mutants, these evolved strains achieved significantly enhanced biomass accumulation relative to the wild-type parent. Whole-genome resequencing across the eight lineages revealed a diverse landscape of novel mutations spanning photosystem assembly, regulatory networks, and metabolic maintenance, demonstrating that cyanobacteria employ distinct polygenic pathways to achieve extreme high-light tolerance.

## Results

### Phenotype characterizations of the ALE strains

Originally, the optimized light density for wild type (WT) of Syn6803 was about 50 μmol photons/m^2^/s, whose growth was gradually inhibited with the increasing light density (**Fig. S1A**). Via nearly two years’ ALE (86 passages) with gradually increased light density (**Fig. 1A** and **1B**), we obtained eight independent strains named from HL-1 to HL-8 that can endure HL up to 750 μmol photons/m^2^/s. As shown in **Fig. 1C**, growth of eight ALE strains under 750 μmol photons/m^2^/s was significantly superior to that of WT. Meanwhile, their final growth was similar to that of WT under normal light of 50 μmol photons/m^2^/s (**Fig. S1B**) though they are lower first three days, suggesting limited growth trade-off happened during the evolution process. Excitingly, we found that these ALE strains could actually withstand light intensities up to 2000 μmol photons/m^2^/s without any further directed evolution (**Fig. 1D**). This suggested that a tolerance level of 750 μmol photons/m^2^/s was sufficient for the strain to cope with even higher light intensities. The visual color of eight ALE strains varied from each other under 750 μmol photons/m^2^/s, suggesting their changed pigment compositions (**Fig. 1E**). Accordingly, the absorption spectrum measured under 750 μmol photons/m^2^/s at the 7^th^ day showed that the main pigments^17^ including chlorophyll, carotenoids, and phycobiliproteins varied among different ALE strains but higher than in WT (**Fig. 1F**). To detailly characterize their differences, we tracked their contents from the 2^nd^ to 7^th^ day. As shown in **Fig. 1G**, compared with that on the 2^nd^ day, content of chlorophyll in WT sharply decreased by 64.57% on the 3^rd^ day and maintained at about 51.57% on the 7^th^ day, which could be a common strategy utilized by Syn6803 to reduce the light absorption. On the contrary, they remained almost stable (71.67% to 98.13%) in all eight ALE strains. Carotenoids are the key pigment to photoprotection^18^, whose content in WT were significantly lower (by 40.75% to 64.90%) than that in eight ALE strains over the culture period (**Fig. 1H)**. Serving as the light capture (450–650 nm) complex, contents of phycobiliproteins decreased gradually in WT (finally decreased by 66.57%) while increased gradually in eight ALE strains (finally increased by 7.09% to 116.86%) (**Fig. 1I**). On one hand, we calculated the relative total content of these pigments and their fractions in eight ALE strains using the 7^th^ day’s data. The differences may partially explain their different colors under HL (**Fig. 1J)**. On the other hand, these preliminary results indicate that photosynthesis of the ALE strains were not significantly altered under HL, as their content of light-harvesting pigments is the same as that of the WT under normal light of 50 μmol photons/m^2^/s (**Fig. S1C**), but is much higher than that of the WT under HL. Similarly, we measured the contents of chlorophyll, carotenoids, and phycobiliproteins under a light density of 2000 μmol photons/m^2^/s on the 2^nd^ and 7^th^ day (**Fig. S1D, S1E**, and **S1F**). All the results suggested eight ALE strains have adapted to the HL conditions and the different phenotype like culture colors indicated they may perform diverse strategies against HL.

**Figure 1.**
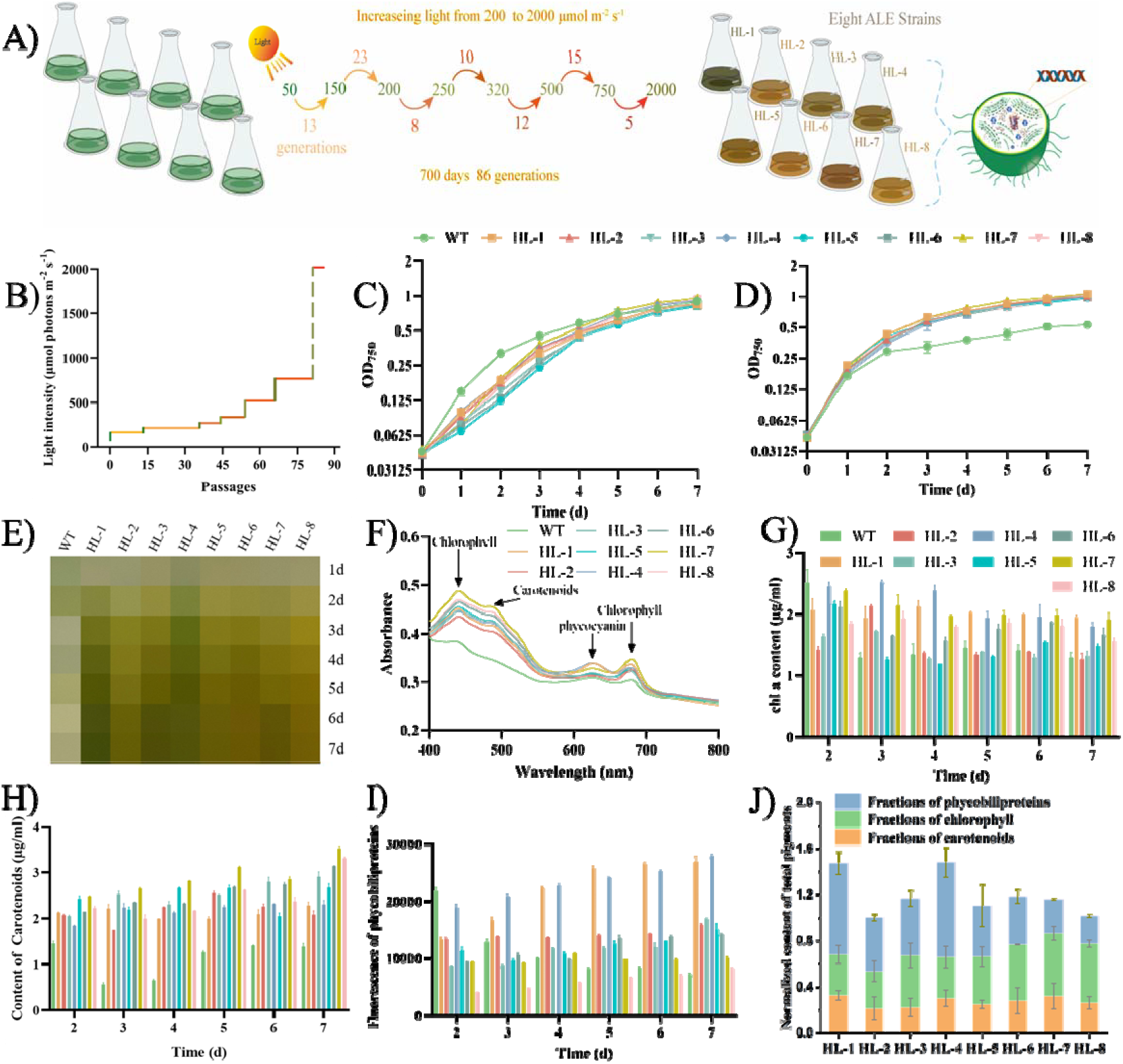
Schematic and phenotype characterizations of the ALE strains. The error bars represented the standard deviation between three biological replicates. **A)** Schematic of the ALE process towards HL tolerance. **B)** The details of increased light density during passages. The growth patterns of WT and eight ALE strains cultivated under **C)** NL (50 μmol photons/m^2^/s) and **D)** HL (750 μmol photons/m^2^/s).**E)** The daily visible color change of WT and eight ALE strains cultivated under HL (750 μmol photons/m^2^/s). **F)** The absorption spectrum of WT and eight ALE strains cultivated under HL (750 μmol photons/m^2^/s) at the 7^th^ d. The daily **G)** chlorophyll, **H)** carotenoids, and **I)** phycobiliproteins change of WT and eight ALE strains cultivated under HL (750 μmol photons/m^2^/s). **J)** The relative content of pigments and their compositions in eight ALE strains cultivated under HL (750 μmol photons/m^2^/s) at the 7^th^ d.

### ALE strains exhibited enhanced photosynthesis under HL condition

Previously, Dann et al. (2021)^15^ investigated reduced biomass accumulation in ALE strains tolerant to HL compared to that of WT under 2000 μmol photons/m^2^/s, suggesting light utilization is not improved with the enhanced HL tolerance. In this study, chlorophyll a fluorescence induction kinetic analysis was used to evaluate the photosynthetic capability of ALE strains under different light conditions. As shown in **Fig. 2A**, the relative ratio of Fv/Fm (reflecting the maximum photochemical quantum yield of PS II) in eight ALE strains were all significantly higher than that in WT, with an increase from 110.10% (HL-6) to 288.89% (HL-8), suggesting enhanced activity of PSII under HL. Consistently, ψ_0_ and Φ_E0_ represented the electron transport capability were significantly higher in eight ALE strains than that in WT (**Fig. 2A**). Meanwhile, DIo/RC indicating the dissipated energy flux per reaction center was lower in eight ALE strains than that in WT (**Fig. 2A**). All these results suggested that ALE strains maintained higher activity of PSII under HL. The enhanced photosynthetic capability of ALE strains was demonstrated via tracking the dissolved oxygen in medium. As shown in **Fig. 2B**, oxygen generation of ALE strains were improved by 4.59% (HL-1) to 19.27% (HL-2) under the tested condition, which was consistent with the results observed in **Fig. 2A**. Considering the enhanced photoreaction of ALE strains under HL, we further investigated the changes of their carbon fixation capability. The content of glycogen, which is the main carbon sink in cyanobacteria, was found significantly increased in all the 8 ALE strains compared to that in WT both under 750 and 2000 μmol photons/m^2^/s (**Fig. 2C**), even higher than that in WT under its optimal growth condition (**Fig. S2A**), suggesting an enhanced carbon fixation in ALE strains. Meanwhile, biomass accumulation in eight ALE strains was not affected under HL compared to that of WT, which increased by 121.71% (HL-1) to 168.36% (HL-2) under 750 μmol photons/m^2^/s and by 88.89% (HL-1) to 123.19% (HL-2) under 2000 μmol photons/m^2^/s (**Fig. 2D**). Notably, biomass accumulation in ALE strains like HL-2 under HL was even higher than that of WT under its optimal condition of 50 μmol photons/m^2^/s (**Fig. S2B**). Via the TEM analysis, we surprisingly found numbers of carboxysomes in HL-2 (exhibiting the most glycogen and biomass accumulation) were higher than that in WT, which may partially explain the enhanced carbon fixation (**Fig. 2E**). A similar phenomenon was observed in the other two randomly selected ALE strains HL-4 and HL-7 also with higher biomass accumulation (**Fig. S2C**). Thus, we believed that enhanced HL utilization and carbon fixation was the main reason that ALE strains maintained low ROS and single oxygen content under HL (both 750 and 2000 μmol photons/m^2^/s; **Fig 2G, S2D** and **S2E**). Encouraged by the above findings, we introduced the sucrose permease encoding gene *cscB* into the ALE strains to evaluate whether they are better chassis for “photosynthetic cell factories” (**Fig. 2H; Table S1**). All strains grew under their optimal condition (50 and 750 μmol photons/m^2^/s for WT and ALE strains, respectively) and salt stimulation was added on the 4^th^ day (**Fig. 2I**). As expected, an increase varied from 29.62% (HL-6-CscB) to 208.33% (HL-2-CscB) in production of sucrose was realized in eight ALE strains compared to that in WT, among which HL-1 showed the highest increase (**Fig. 2J**). Under normal conditions, both growth and production of sucrose in eight ALE strains were comparable to that of WT (**Fig. S2F** and **S2G**). All these results suggested inherent changes appeared in ALE strains, making them a better chassis for HL utilization and CO_2_ fixation.

**Figure 2.**
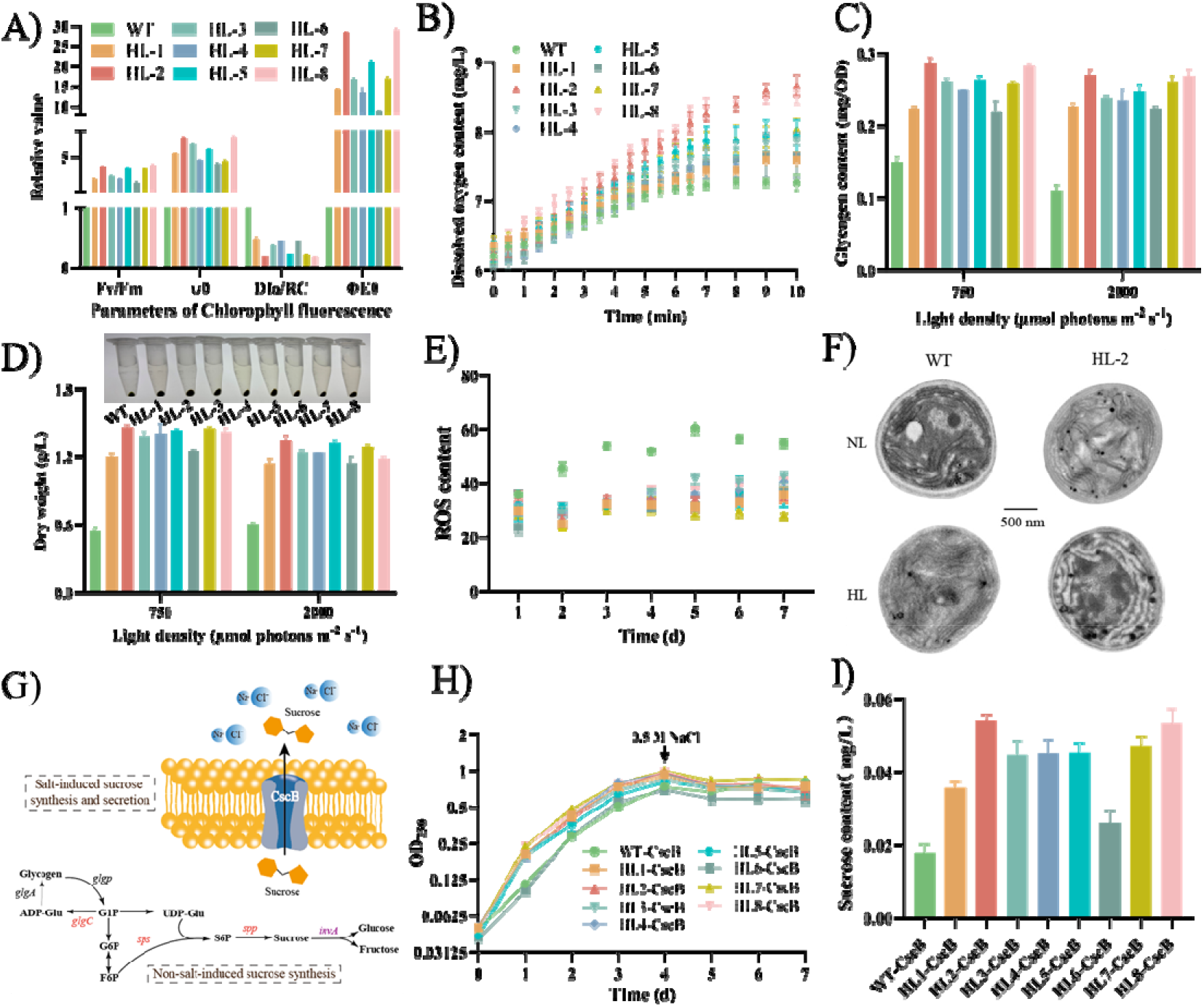
Photosynthesis and carbon fixation capability evaluation of eight ALE strains. The error bars represented the standard deviation between three biological replicates. **A)** Results of chlorophyll fluorescence parameters and **B)** Oxygen evolution of WT and eight ALE strains cultivated under HL (750 μmol photons/m^2^/s). **C)** Glycogen and **D)** biomass accumulation of WT and eight ALE strains cultivated under HL (750 and 2000 μmol photons/m^2^/s). **E)** ROS content of WT and eight ALE strains cultivated under HL (750 μmol photons/m^2^/s). **F)** TEM analysis of WT and HL-2 under NL (50 μmol photons/m^2^/s) and HL (750 μmol photons/m^2^/s). **G)** Schematic of sucrose synthesis and secretion pathways. **H)** Growth patterns and **I)** sucrose production in constructed strains cultivated under NL (50 μmol photons/m^2^/s) for WT-CscB and HL (750 μmol photons/m^2^/s) for HL-1-CscB…, HL-8-CscB.

### Genome re-sequencing identified key genes responding to HL tolerance in ALE strains

To explore the mechanisms against HL, the genome sequences of eight ALE strains were re-analyzed to find the mutations. Totally, 77 mutations including 44 substitution mutations, 8 deletion mutations, and 24 insertion mutations, were identified in eight ALE strains through the genome sequencing, 63 of which involving 57 genes were demonstrated via Sanger sequencing (**Table S2**). Among them, mutations in *slr0193, slr0320, sll0665, rpoC2, slr0758, spoT, slr1862* and *rpoB* were shared in at least two ALE strains (**Fig. 3A**). In addition, up to 38 unique mutations were found in HL-6 while several unique mutations were respectively found in the other seven strains (**Fig. 3A**). All the identified genes except 4 transposon encoding genes and *slr1512* (whose relationship with HL tolerance have been studied in our previous work^9^) were respectively deleted from the genome of WT, leading to 52 gene-knockout strains (**Fig. 4B, Table S1**). Via comparing the growth patterns of WT and gene-knockout strains under 750 μmol photons/m^2^/s, 21 of 52 genes like *slr0320, slr0193, slr0758, glgX, slr1147*, and *slr1513* were found significantly related with HL response (**Fig. 3B, S3A-G**). Among them, deletion of *slr0320* (Δ*slr0320*) encoding a SAM-domain protein decreased the HL tolerance of WT but deletion of *slr0193* (Δ*slr0193*) encoding an RNA-binding protein enhanced the tolerance (**Fig. 3B**). Notably, mutations of *slr0320* and *slr0193* co-occurred in HL-5 and HL-7 as well as HL-1, HL-3, and HL-6 (**Fig. 3A**), respectively. Among them, deletion of slr0320 encoding a SAM-domain protein decreased the high-light tolerance of WT, whereas deletion of slr0193 encoding an RNA-binding protein enhanced tolerance (Fig. 3B). Notably, mutations in slr0320 and slr0193 co-occurred in multiple ALE strains (e.g., HL-5, HL-7, HL-1, HL-3, and HL-6; **Fig. 3A**). For other mutations, we briefly explored the roles of *slr1513* (*sbtB*) as it specifically regulated *slr1512*, which has been demonstrated related with HL tolerance^9^. In this study, mutation of Slr1513 (Y87C) was found in HL-7 and HL-8. As the structure of both Slr1513 and Slr1512 have been elucidated^19, 20^, we analyzed their interactions via molecular docking. As shown in **Fig. 3C**, we found the mutation of Slr1513 just existed in its interaction region with Slr1512, decreasing its regulation on Slr1512. Consistent with the previous study, increased content of ATP under HL would weaken the regulation of Slr1513 on Slr1512, which could further enhance the carbon uptake of Slr1512. In this study, we speculated that altered interaction between Slr1513 and Slr1512 may decrease the uptake of HCO ^-^ as we have demonstrated that decreased HCO_3_ ^-^ would restrict the electron transfer in PSII thus relieving the stress from utilizing excess photons under HL^9^. Interestingly, deletion of *slr0758* encoding KaiC1 enhanced the tolerance to HL. Slr0758 exhibited three kinds of mutations including A329P, T178M, and Q153U in HL-2, HL-5, and HL-6, respectively. The A329 mutation reduces inter-residue interactions. The proline residue introduced by this mutation forms a cyclic structure between its side chain and the backbone amino group, thereby restricting conformational flexibility. Due to its unique pyrrolidine ring, proline often plays a special role in protein structures and may disrupt α-helical regions. This disruption occurs by breaking the hydrogen bond network essential for maintaining secondary structures, potentially introducing a helix-breaking effect and impairing enzymatic activity (**Fig. 3D**). The T178M mutation replaces threonine—which normally participates in hydrogen bond formation—with methionine, which lacks this capacity for interaction. This substitution may alter the local structure, such as α-helices or β-sheets, and consequently affect protein function (**Fig. 3D**). The Q153U mutation results in a premature termination of translation, leading to a truncated protein. This shortened form is likely to undergo structural changes, potentially resulting in a complete loss of function. We believe the mutation of Slr0758 benefited the HL tolerance due to two reasons: i) Several studies have demonstrated the rhythmic system composed of KaiABC also regulated the gene expression under constant light except of rhythmic light^21, 22^; ii) halotolerant cyanobacteria *Halothece* sp. PCC7418 was found to prioritize the adaptation to salt stress by attenuation of circadian rhythmicity^23^.

**Figure 3.**
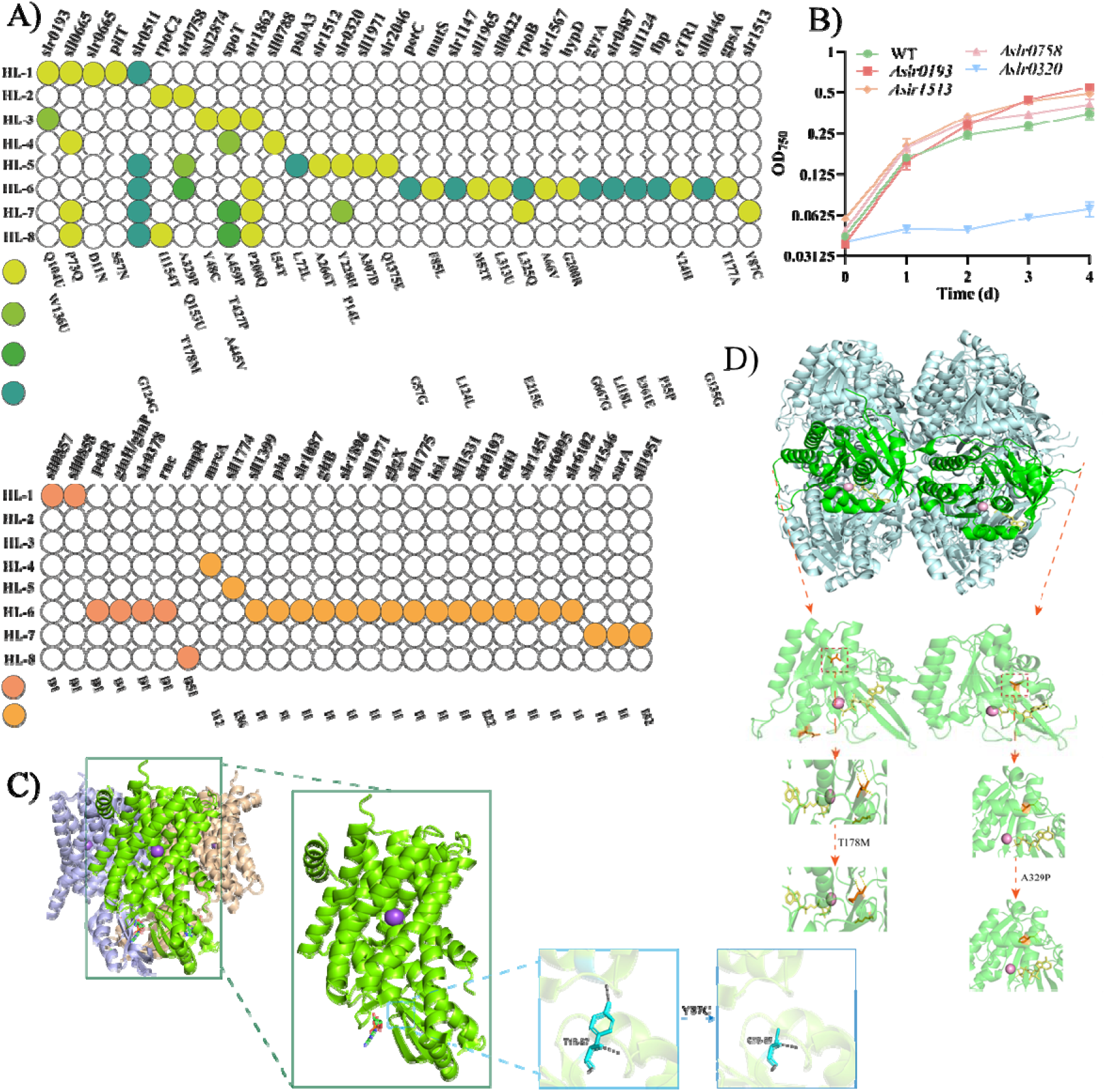
Genome re-sequencing results and key genes responding to HL tolerance in eight ALE strains. The error bars represented the standard deviation between three biological replicates. **A)** Summary of mutation sites identified in the eight ALE strains. **B)** Growth patterns of WT, Δ*slr0758*, Δ*slr0193*, Δ*slr0320*, and Δ*slr1513* cultivated under HL (750 μmol photons/m^2^/s). **C)** Predicted structural changes in the interaction between Slr1512 and Slr1513 after mutation. **D)** Predicted structural changes of Slr0758 protein after mutation.

## Discussion

ALE has proven to be a powerful approach not only for obtaining stress-tolerant strains, providing optimized chassis candidates for synthetic biology in cyanobacteria. In previous studies, HL-tolerant Syn6803 strains were obtained through different ALE strategies^15, 16^. Similarly, once strains acquired HL tolerance, they often exhibited altered pigmentation (non-blue-green coloration) and slower growth under low-light conditions compared to the wild type. Differently, in this study, the biomass accumulation of eight HL-tolerant strains was not impaired but significantly improved. This suggests that these strains did not merely acquire mechanisms to cope with photodamage but also enhanced their ability to utilize HL more efficiently. Among these, HL-2, HL-5, HL-7, and HL-8 demonstrated markedly enhanced maximum quantum yield, oxygen evolution rates, and carbon fixation capacities. Furthermore, when tested for heterologous product yields, similar improvements were observed, indicating their potential as advanced chassis strains. HL-2 stood out as the most promising strain due to its superior performance in biomass accumulation, sucrose and glycogen production, and the highest quantum yield. Our findings also support the conclusions of Ducat et al.^24^, which proposed that sucrose biosynthesis can serve as a carbon sink under HL conditions, aiding in HL acclimation. Previous systems biology analyses have shown that HL stress triggers multiple response mechanisms in wild-type cyanobacteria, such as reducing light harvesting, quenching excess energy, and inducing alternative electron transport pathways^2^. This diversity in response strategies implies that multiple routes can lead to HL tolerance. Genetic analyses in evolved strains from earlier studies identified key mutations in genes such as *slr0844, sll1098, hik26*, and *slr1916*^15, 16^.

Despite these advances, several directions remain open for future investigation. First, while we identified distinct mutations at the genomic level among independently evolved strains, integrated omics approaches—including transcriptomics and metabolomics—could help map out strain-specific metabolic networks under HL conditions, revealing diverse adaptation routes. Potential synergistic interactions among these genes also warrant further study. Lastly, our photosynthetic measurements revealed functional differences in carbon fixation and photosynthesis among HL strains and between HL strains and wild-type. Preliminary TEM analysis suggests possible variation in carboxysome structure and size, hinting at structural changes in photosynthetic machinery. Structural biology studies of key photosynthetic complexes and carboxysomes may provide deeper insights into their adaptations. In conclusion, we successfully obtained eight HL-tolerant strains through independent directed evolution and demonstrated that these strains adopt distinct strategies to cope with HL stress. Our findings contribute novel perspectives to the understanding of HL tolerance mechanisms in photosynthetic microorganisms and offer valuable genetic resources for synthetic biology applications.

## Methods

### Strains, culture conditions and growth patterns

The WT, as well as the Syn6803 strains constructed and evolved in this study, were cultured in standard BG11 medium (pH 7.5) under an atmospheric CO_2_ concentration in an HNY-211B Illuminating Shaker at 30°C and 130 rpm (Honour, Tianjin, China). The incubator was equipped with a set of LED lights capable of adjusting the light intensity between 50 and 2000 µmol photons/m^2^/s. The cell density was determined at OD_750 nm_ using an ELx808 Absorbance Microplate Reader (BioTek, VT, USA). For growth experiments, 5 mL of fresh cells at OD_750 nm_ of 0.2 were collected by centrifugation and inoculated into 20 mL of BG11 liquid medium in a flask. The WT and the desired strain(s) were then cultured under specific light intensities as required. Every 24 hours, 200 μL of culture samples were taken and measured at OD_750 nm_. Growth experiments were repeated at least six times to confirm the growth patterns.

### ALE strategy

The WT was initially inoculated into 8 shaking flasks with an initial OD_750_ nm of 0.04. These 8 replicates were cultured under a light intensity of 100 µmol photons/m^2^/s and passaged once they reached their highest cell density, which typically took around 7-10 days. The light intensity was gradually increased (100, 150, 200, 350, 600, 750 and 2000 µmol photons/m^2^/s) until the 8 replicates could reach a similar cell density as that achieved under normal light conditions (50 µmol photons/m^2^/s). Throughout the ALE process, the 8 replicates gradually adapted to increasing light intensities, which ranged from 100 to 2000 µmol photons/m^2^/s. Finally, to analyze the ALE strain in the following studies, 8 single colonies were separated from the 8 according ALE strains (named from HLT-1to HLT-8), respectively.

### Chlorophyll a and carotenoids quantitation

To determine the content of chlorophyll a (Chl a) and carotenoids, we collected samples of both the WT and ALE strains from HL-treated cultures (750 or 2000 µmol photons/m^2^/s). Cells with a volume*OD_750 nm_ of 0.5 were centrifuged at 12000 x g for 2 minutes at 4°C to obtain pellets. The pellets were then re-suspended in 1 mL of pre-cooled (4°C) methanol and incubated at 4°C for 20 minutes to extract the pigments. The supernatant was collected by centrifugation at 12,000 x g and 4°C for 10 minutes. Finally, the supernatant was measured at 720 nm, 665 nm, and 470 nm. To calculate the content of Chl a, we used the formula: Chl a [µg/ml] = dilution ratio *12.9447*(A_665_ -A_720_)^25^. For the carotenoid content, we used the formula: Carotenoids [µg/ml] = dilution dilution ratio *[1,000*(A_470_-A_720_) -2.86*(content of Chl a)]/221^26^.

### ROS and singlet oxygen measurement

For ROS and singlet oxygen measurement, cell samples (volume*OD_750 nm_=1) including HL-treated (750 or 2000 µmol photons/m^2^/s) WT and ALE strains were harvested and resuspended with BG11. To measure ROS content, we used a Reactive Oxygen Species Assay Kit (Beyotime, Shanghai, China) and followed the kit’s protocol. Briefly, the non-fluorescent probe DCFH-DA could be oxidized by intracellular ROS into fluorescent DCF, which could be detected at EX 488 nm and EM 525 nm using an F-2700 fluorescence spectrophotometer (Hitachi, Tokyo, Japan). To measure singlet oxygen content, we used the SOSG singlet oxygen fluorescence probe (Meilunstar, DaLian, China) and followed a published protocol. The SOSG stained sample was detected using the same fluorescence spectrophotometer at EX 504 nm and EM 525 nm.

### Absorption spectra and chlorophyll fluorescence parameters determination

Cell samples from both HL-treated cultures (750 or 2000 µmol photons/m^2^/s) of WT and ALE strains were collected, followed by normalization using OD_750 nm_ and subsequent measurement using the UV-1601 spectrophotometer (Beifen-Ruili, Beijing, China). The obtained data were analyzed using UVprobe software (Version X). Additionally, fluorescence parameters of chlorophyll, including Fv/Fm, OJIP curve, and other related chlorophyll fluorescence parameters, were measured using the AquaPen AP 110/C Handheld PAM Fluorometer (FluorCam, Drásov, Czech Republic) for HL-treated (750 or 2000 µmol photons/m^2^/s) WT and ALE strains.

### Cell dry weight and glycogen measurement

For measurement of cell dry weight, 20 mL samples of both WT and ALE strains were obtained by centrifugation (7000 x g; 10 minutes; 4°C) after 7 days of cultivation. The resulting cell pellets were then transferred to pre-weighed 1.5 mL centrifuge tubes and immediately frozen in liquid nitrogen. The samples were subsequently dried using an LGJ-10 Vacuum Freeze Drier (Songyuan Huaxing, Beijing, China) and re-weighed.

To quantify glycogen, an anthrone method-based kit (Solarbio, Beijing, China) was used. Briefly, both WT and ALE strains (volume*OD750 nm=0.5) were harvested by centrifugation (15000 x g; 2 minutes; 4°C) after 7 days of cultivation. The resulting cell pellets were crushed using a HX-21G Rapid tissue breaking apparatus (Honour, Tianjin, China). The assays were performed according to the protocols provided and the absorbance was measured at OD_620_. A group of standards (0.0125, 0.025, 0.05, 0.1, 0.15, 0.2 g/L) were also measured to obtain the standard curve.

### Genome re-sequencing and transcriptome analysis

The genome sequences of both the WT and 8 ALE strains were obtained using the IlluminaHiseq PE150 platform (Novogene, Beijing, China). The low-quality data were filtered out and mapped to the reference genome of Syn6803 (https://www.ncbi.nlm.nih.gov/nuccore/NC_000911) before the mutation sites were validated via Sanger sequencing (GENEWIZ, Suzhou, China). For transcriptome sequencing, three biological replicates of the WT strain grown under 50 µmol photons/m^2^/s, as well as three replicates of the WT, HLT-2, HLT-4, and HLT-7 strains grown under 750 µmol photons/m^2^/s were collected after 3 days of cultivation, totaling 15 samples. After centrifugation at 4°C following 4 days of cultivation, the samples were immediately frozen in liquid nitrogen for RNA-seq (GENEWIZ, Suzhou, China). To analyze the data, the raw reads were filtered to obtain clean reads by removing low-quality reads, adapter contamination, and reads with unknown bases (N reads). The clean data were then mapped to the reference genome of Syn6803 (https://www.ncbi.nlm.nih.gov/nuccore/NC_000911), and the reads counts were normalized to the aligned FPKM (Fragments Per Kilobase of transcript, per Million mapped reads) to obtain the relative expression levels. Differential expression analysis was performed using DESeq2 (V1.6.3) to compare the samples, with genes having a fold change ≥2 and an adjusted p-value ≤0.05 being considered differentially expressed genes.

### Gene deletion, complementation, replacement, site mutation and overexpression

The genetic modification of Syn6803 was accomplished using natural transformation through homologous recombination. To delete a gene, a chloramphenicol-resistant cassette was used to replace the target gene. To complement or replace a gene, the original gene fragment was fused to a spectinomycin-resistant cassette, along with two homologous arms, and then introduced into the gene-knockout strain. To overexpress a gene, the strong promoter P_cpc560_ was used to drive the expression of the target gene. The fragment was fused with a chloramphenicol-resistant cassette and two homologous arms, and then integrated into the neutral site *slr0168*^27^. The constructed strains were screened on agar plates using antibiotics (10 mg/L chloramphenicol or spectinomycin), and then validated via colony PCR and Sanger sequencing.

### Sucrose-producing strain construction and sucrose quantitation

The sucrose permease encoding gene, *cscB*, was utilized to secrete accumulated sucrose under salt stress^28^. The construction strategy was similar to the gene overexpression method mentioned above. Sucrose quantitation was performed using a Sucrose Assay Kit (Megazyme, Wicklow, Kingdom of Ireland) following the protocol. The constructed strains were cultured under 50 or 750 µmol photons/m^2^/s and induced with 0.5 M NaCl on the fourth day. Samples were collected from the supernatant of 1 mL cultures via centrifugation (15000 x g; 2 minutes; 4°C).

### Protein structure prediction and molecular docking

The amino acid sequence (FASTA format) of the target protein was submitted to ColabFold (based on AlphaFold) for 3D structure prediction. Predicted PDB structures were visualized and analyzed using PyMOL.

### Phycocyanin measurement

Cells were cultured until OD_750_ ≈ 0.020. Cells were harvested by centrifugation at 8000 rpm for 5 min, and the supernatant was discarded. The pellet was resuspended in 200 μL sterile water and incubated in a dark 96-well plate for 30 minutes. The absorbance ratio at 630 nm and 660 nm was measured to estimate phycocyanin content.

### Significance analysis

Student’s t test was used to evaluate the data significance.

### Accession numbers

The GenBank reference sequences used for the genome assembly and transcriptome mapping of WT and ALE strains were NC_000911.1 (chromosome), AP004311 (Kazusa, pSYSA), AP004310 (Kazusa, pSYSM), AP004312 (Kazusa, pSYSG), AP006585 (Kazusa, pSYSX), CP003270 (Freiburg, pCA) and CP003271 (Freiburg, pCB). All the re-sequenced genomes in this study have been deposited at Sequence Read Archive (SRA) of NCBI (Accession number PRJNA1297135). All the re-sequenced genomes in this study have been deposited at SRA (Accession number RJNA1297135)

## Data availability

The genome sequencing and transcriptomic data have been uploaded to SRA of NCBI. Other data supporting the findings of this study are available within the paper and its Supplementary Information.

## Acknowledgements

This research was supported by grants from the National Key Research and Development Program of China (Grant no. 2024YFA0919700), the National Natural Science Foundation of China (Grant nos. 32371486, and 32270091), and the Haihe Laboratory of Sustainable Chemical Transformations.

## Author contributions

Tao Sun was responsible for the entire experimental design, data analysis, and drafting and reviewing of the paper. Kungang Pan and Yaru Xie performed the main experiment and reviewed the draft. Shubin Li and Congzhuang Li performed some experiments related to phenotype identification. Dainlin Liu and Xiaofei Zhu was involved in experimental design related to ALE strain analysis. Weiwen Zhang and Lei Chen provided supervision, reviewed drafts, and edited the paper.

## Additional information

Supplementary data to this article can be found online.

