## Supplemental Figures and Tables for "Engineering light robustness: Adaptive evolution uncovers new genetic determinants of HL tolerance in *Synechocystis*"

**Adaptive evolution uncovers a conserved post-transcriptional module governing photosynthetic robustness**

*^1^ School of Synthetic Biology and Biomanufacturing, Tianjin University, Tianjin, China; ^2^ Frontier Science Center for Synthetic Biology & Key Laboratory of Systems Bioengineering, Ministry of Education of China, Tianjin 300072, P.R. China; ^3^* *State Key Laboratory of Synthetic Biology, Tianjin University, Tianjin, 300072, PR China;  ^4^ Center for Biosafety Research and Strategy, Tianjin University, Tianjin 300072, P.R. China; ^5^ Haihe Laboratory of Sustainable Chemical Transformations, Tianjin 300192, China.*

\# These three authors contributed equally to the paper.

* To whom all correspondence should be addressed:

Prof. Dr. Weiwen Zhang

Laboratory of Synthetic Microbiology

School of Chemical Engineering & Technology

Tianjin University, Tianjin 300072, P.R. China

Prof. Dr. Lei Chen

Laboratory of Synthetic Microbiology

School of Chemical Engineering & Technology

Tianjin University, Tianjin 300072, P.R. China

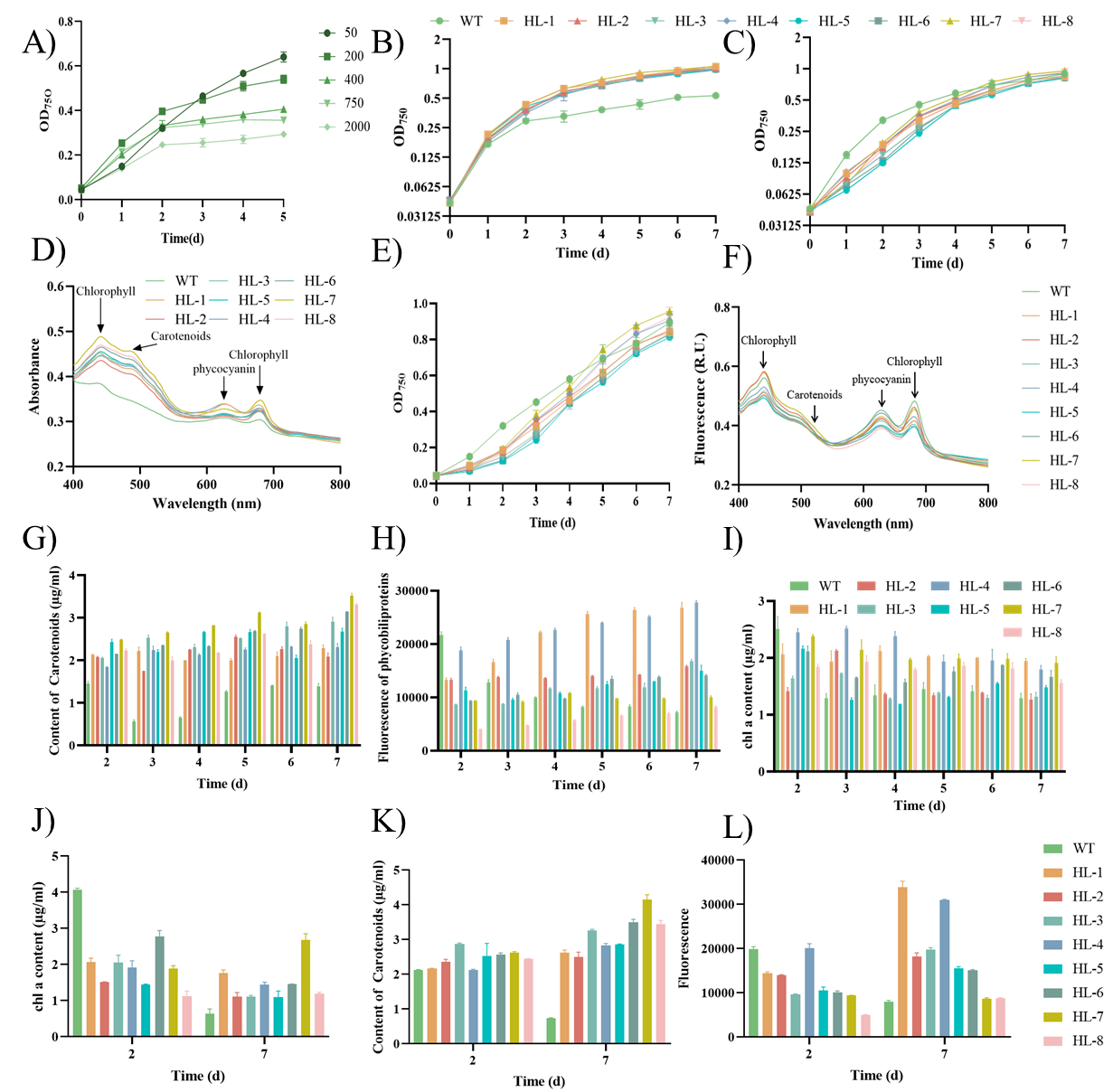

**Figure S1. A)** Growth curves of WT under different light density. The growth patterns of WT and eight ALE strains cultivated under **B)** HL (750 μmol photons/m^2^/s) and **C)** NL (50 μmol photons/m^2^/s). **D)** The absorption spectrum of WT and eight ALE strains cultivated under HL (750 μmol photons/m^2^/s) at the 7^th^ d. **E)** Growth patterns and **F)** absorption spectrum of WT and eight ALE strains cultivated under NL (50 μmol photons/m^2^/s). The daily **G)** chlorophyll, **H)** carotenoids, and **I)** phycobiliproteins change of WT and eight ALE strains cultivated under HL (750 μmol photons/m2/s). J) The relative content of pigments and their compositions in eight ALE strains cultivated under HL (750 μmol photons/m^2^/s) at the 7^th^ d. The content of **J)** chlorophyll, **K)** carotenoids, and **L)** phycobiliproteins change of WT and eight ALE strains cultivated under HL (2000 μmol photons/m^2^/s) at the 2^nd^ and 7^th^ d.

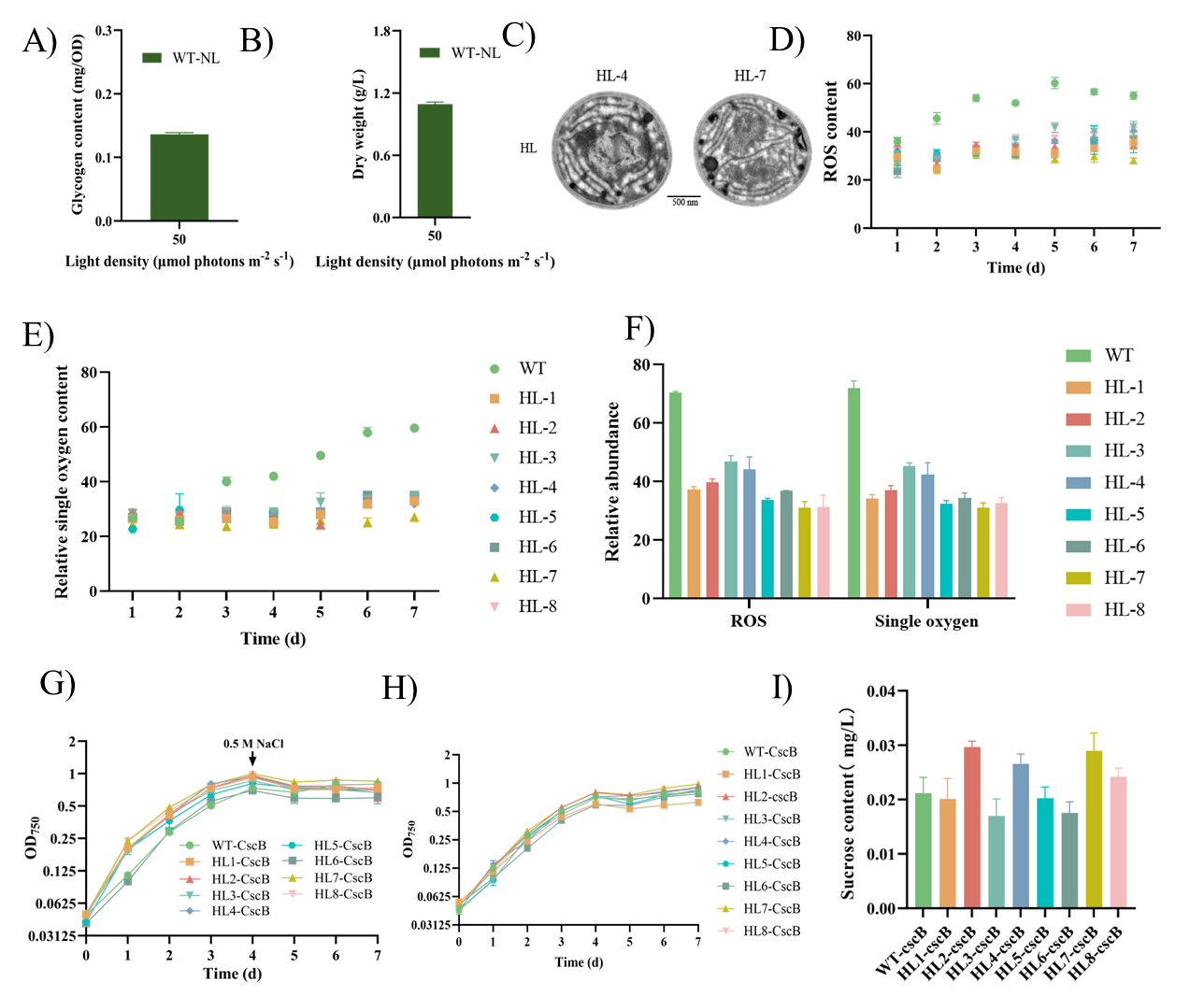

**Figure S2.** The **A)** glycogen and **B)** biomass accumulation of WT under NL (50 μmol photons/m^2^/s). **C)** TEM image of HL-4 and HL-7. **D)** ROS content of WT and eight ALE strains cultivated under HL (750 μmol photons/m^2^/s)**. E)** The daily relative single oxygen content change of WT and eight ALE strains cultivated under HL (750 μmol photons/m^2^/s). **F)** The relative ROS and single oxygen content change of WT and eight ALE strains cultivated under HL (2000 μmol photons/m^2^/s) at the 7^th^ d. **G)** Growth patterns in constructed strains cultivated under NL (50 μmol photons/m^2^/s) for WT-CscB and HL (750 μmol photons/m^2^/s) for HL-1-CscB…, HL-8-CscB. **H)** Growth patterns and **I)** sucrose production in constructed strains cultivated under NL (50 μmol photons/m^2^/s).

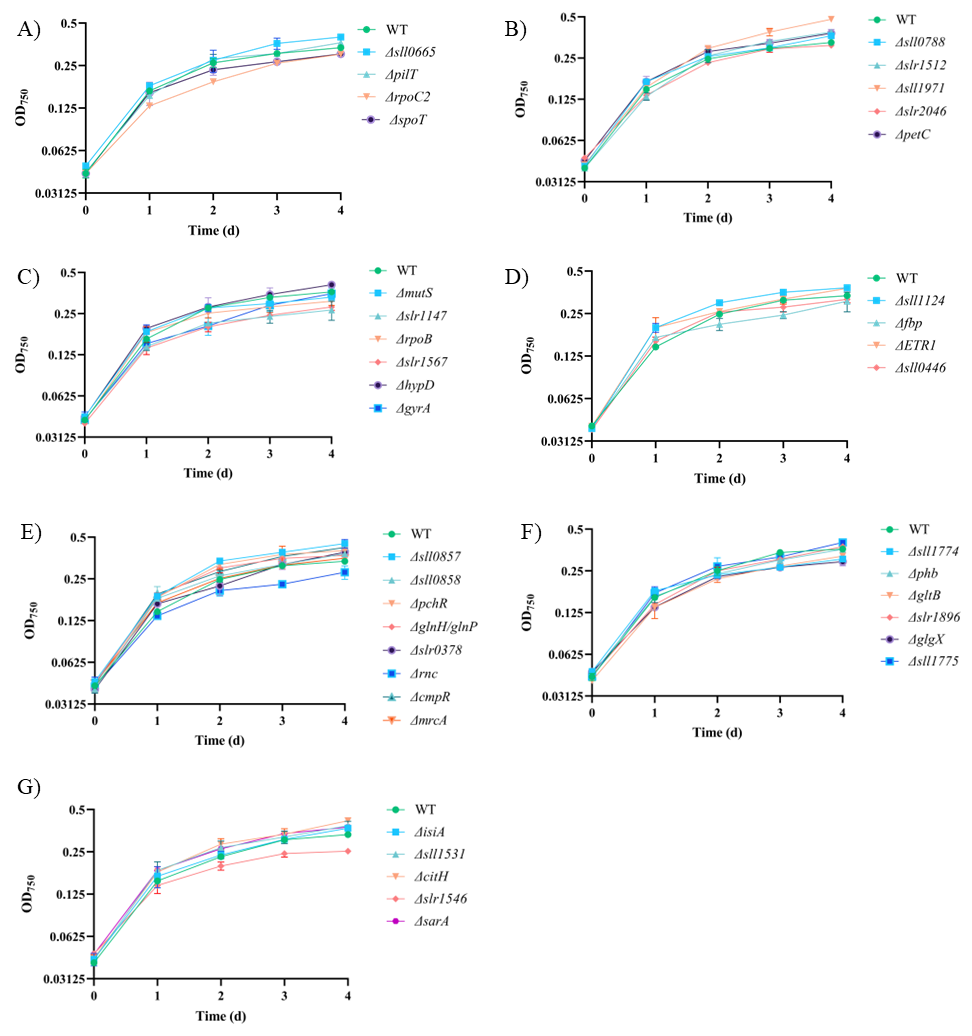

**Figure S3. A)-G)** Growth patterns of WT and gene deleted strains cultivated under HL (750 μmol photons/m^2^/s).

Table S1. Strains constructed and used in this study.

| Strain Name | Descriptions |
| --- | --- |
| HL-1 | Through approximately 2 years of adaptive laboratory evolution, an independent strain named HL-1 was obtained that tolerated 2000 μmol photons/m^2^/s HL |
| HL-2 | Through approximately 2 years of adaptive laboratory evolution, an independent strain named HL-2 was obtained that tolerated 2000 μmol photons/m^2^/s HL |
| HL-3 | Through approximately 2 years of adaptive laboratory evolution, an independent strain named HL-3 was obtained that tolerated 2000 μmol photons/m^2^/s HL |
| HL-4 | Through approximately 2 years of adaptive laboratory evolution, an independent strain named HL-4 was obtained that tolerated 2000 μmol photons/m^2^/s HL |
| HL-5 | Through approximately 2 years of adaptive laboratory evolution, an independent strain named HL-5 was obtained that tolerated 2000 μmol photons/m^2^/s HL |
| HL-6 | Through approximately 2 years of adaptive laboratory evolution, an independent strain named HL-6 was obtained that tolerated 2000 μmol photons/m^2^/s HL |
| HL-7 | Through approximately 2 years of adaptive laboratory evolution, an independent strain named HL-7 was obtained that tolerated 2000 μmol photons/m^2^/s HL |
| HL-8 | Through approximately 2 years of adaptive laboratory evolution, an independent strain named HL-8 was obtained that tolerated 2000 μmol photons/m^2^/s HL |
| WT-CSCB | Introduction of the sucrose permease encoding gene *cscB* in WT strain |
| HL1-CSCB | Introduction of the sucrose permease encoding gene *cscB* in ALE HL-1 strain |
| HL2-CSCB | Introduction of the sucrose permease encoding gene *cscB* in ALE HL-2 strain |
| HL3-CSCB | Introduction of the sucrose permease encoding gene *cscB* in ALE HL-3 strain |
| HL4-CSCB | Introduction of the sucrose permease encoding gene *cscB* in ALE HL-4 strain |
| HL5-CSCB | Introduction of the sucrose permease encoding gene *cscB* in ALE HL-5 strain |
| HL6-CSCB | Introduction of the sucrose permease encoding gene *cscB* in ALE HL-6 strain |
| HL7-CSCB | Introduction of the sucrose permease encoding gene *cscB* in ALE HL-7 strain |
| HL8-CSCB | Introduction of the sucrose permease encoding gene *cscB* in ALE HL-8 strain |
| Δslr0193 | Replacement of in situ gene *slr0193* in WT for Cm^R^ resistance |
| Δsll0665 | Replacement of in situ gene *sll0665* in WT for Cm^R^ resistance |
| ΔpilT | Replacement of in situ gene *pilT* in WT for Cm^R^ resistance |
| ΔrpoC2 | Replacement of in situ gene *rpoC2* in WT for Cm^R^ resistance |
| Δslr0758 | Replacement of in situ gene *slr0758* in WT for Cm^R^ resistance |
| ΔspoT | Replacement of in situ gene *spoT* in WT for Cm^R^ resistance |
| Δsll0788 | Replacement of in situ gene *sll0788* in WT for Cm^R^ resistance |
| Δslr1512 | Replacement of in situ gene *slr1512* in WT for Cm^R^ resistance |
| Δslr0320 | Replacement of in situ gene *slr0320* in WT for Cm^R^ resistance |
| Δsll1971 | Replacement of in situ gene *sll1971* in WT for Cm^R^ resistance |
| Δslr2046 | Replacement of in situ gene *slr2046* in WT for Cm^R^ resistance |
| ΔpetC | Replacement of in situ gene *petC* in WT for Cm^R^ resistance |
| ΔmutS | Replacement of in situ gene *mutS* in WT for Cm^R^ resistance |
| Δslr1147 | Replacement of in situ gene *slr1147* in WT for Cm^R^ resistance |
| ΔrpoB | Replacement of in situ gene *rpoB* in WT for Cm^R^ resistance |
| Δslr1567 | Replacement of in situ gene *slr1567* in WT for Cm^R^ resistance |
| ΔhypD | Replacement of in situ gene *hypD* in WT for Cm^R^ resistance |
| ΔgyrA | Replacement of in situ gene *gyrA* in WT for Cm^R^ resistance |
| Δsll1124 | Replacement of in situ gene *sll1124* in WT for Cm^R^ resistance |
| Δfbp | Replacement of in situ gene *fbp* in WT for Cm^R^ resistance |
| ΔETR1 | Replacement of in situ gene *ETR1* in WT for Cm^R^ resistance |
| Δsll0446 | Replacement of in situ gene *sll0446* in WT for Cm^R^ resistance |
| Δslr1515 | Replacement of in situ gene *slr1515* in WT for Cm^R^ resistance |
| Δslr1522 | Replacement of in situ gene *slr1522* in WT for Cm^R^ resistance |
| Δslr1513 | Replacement of in situ gene *slr1513* in WT for Cm^R^ resistance |
| Δsll1201 | Replacement of in situ gene *sll1201* in WT for Cm^R^ resistance |
| Δsll0857 | Replacement of in situ gene *sll0857* in WT for Cm^R^ resistance |
| Δsll0858 | Replacement of in situ gene *sll0858* in WT for Cm^R^ resistance |
| ΔpchR | Replacement of in situ gene *pchR* in WT for Cm^R^ resistance |
| ΔglnH/glnP | Replacement of in situ gene *glnH/glnP* in WT for Cm^R^ resistance |
| Δslr0378 | Replacement of in situ gene *slr0378* in WT for Cm^R^ resistance |
| Δrnc | Replacement of in situ gene *rnc* in WT for Cm^R^ resistance |
| ΔcmpR | Replacement of in situ gene *cmpR* in WT for Cm^R^ resistance |
| ΔmrcA | Replacement of in situ gene *mrcA* in WT for Cm^R^ resistance |
| Δsll1774 | Replacement of in situ gene *sll1774* in WT for Cm^R^ resistance |
| Δphb | Replacement of in situ gene *phb* in WT for Cm^R^ resistance |
| ΔgltB | Replacement of in situ gene *gltB* in WT for Cm^R^ resistance |
| Δslr1896 | Replacement of in situ gene *slr1896* in WT for Cm^R^ resistance |
| ΔglgX | Replacement of in situ gene *glgX* in WT for Cm^R^ resistance |
| Δsll1775 | Replacement of in situ gene *sll1775* in WT for Cm^R^ resistance |
| ΔisiA | Replacement of in situ gene *isiA* in WT for Cm^R^ resistance |
| Δsll1531 | Replacement of in situ gene *sll1531* in WT for Cm^R^ resistance |
| ΔcitH | Replacement of in situ gene *citH* in WT for Cm^R^ resistance |
| Δslr1546 | Replacement of in situ gene *slr1546* in WT for Cm^R^ resistance |
| ΔsarA | Replacement of in situ gene *sarA* in WT for Cm^R^ resistance |

**Table S2. Identified mutations in eight ALE strains.**

| Gene Name | Mutant strains | Mutation Description |
| --- | --- | --- |
| slr0193 | HL-1/HL-3 | Q104U/W136U |
| sll0665 | HL-1 | D11N |
| sll0665 | HL-1/HL-4/HL-7/HL-8 | P73Q |
| pilT | HL-1 | S57N |
| slr0511 | HL-1/HL-5/HL-6/HL-7/HL-8 | G124G |
| rpoC2 | HL-2/HL-8 | I1154T |
| slr0758 | HL-2/HL-5/HL-6 | A324P/Q153U/T178M |
| ssl2874 | HL-3 | Y48C |
| spoT | HL-3/HL-4/HL-7/HL-8 | A459P/T427P/A445V/A445V |
| slr1862 | HL-3/HL-6/HL-7/HL-8 | P200Q |
| sll0788 | HL-4 | I54T |
| psbA3 | HL-5 | L72L |
| slr1512 | HL-5 | A266T |
| slr0320 | HL-5/HL-7 | Y228H/P14L |
| sll1971 | HL-5 | A316D |
| slr2046 | HL-5 | Q1375E |
| petC | HL-6 | G57G |
| mutS | HL-6 | F85L |
| slr1147 | HL-6 | L124L |
| sll1965 | HL-6 | M98T |
| sll0422 | HL-6 | L313U |
| rpoB | HL-6/HL-7 | E215E/L325Q |
| slr1567 | HL-6 | A66V |
| hypD | HL-6 | G200R |
| gyrA | HL-6 | T690A |
| slr0487 | HL-6 | L118L |
| sll1124 | HL-6 | E961E |
| fbp | HL-6 | P35P |
| ETR1 | HL-6 | Y24H |
| sll0446 | HL-6 | G135G |
| gpsA | HL-6 | T177A |
| slr1513 | HL-7 | Y87C |
| sll0857 | HL-1 | D1 |
| sll0858 | HL-1 | D1 |
| pchR | HL-6 | D1 |
| glnH/glnP | HL-6 | D1 |
| slr0378 | HL-6 | D1 |
| rnc | HL-6 | D1 |
| cmpR | HL-8 | D51 |
| mrcA | HL-4 | I12 |
| sll1774 | HL-5 | I36 |
| sll1399 | HL-6 | I1 |
| phb | HL-6 | I1 |
| slr1087 | HL-6 | I1 |
| gltB | HL-6 | I1 |
| slr1896 | HL-6 | I1 |
| sll1971 | HL-6 | I1 |
| glgX | HL-6 | I1 |
| sll1775 | HL-6 | I1 |
| isiA | HL-6 | I1 |
| sll1531 | HL-6 | I1 |
| slr0193 | HL-6 | I22 |
| citH | HL-6 | I1 |
| slr1451 | HL-6 | I1 |
| slr6102 | HL-6 | I1 |
| slr1546 | HL-7 | I1 |
| sarA | HL-7 | I1 |
| sll1951 | HL-7 | I32 |
